# Direct-print three-dimensional electrodes for large-scale, high-density, and customizable neural interfaces

**DOI:** 10.1101/2023.05.30.542925

**Authors:** Pingyu Wang, Eric G. Wu, Hasan Uluşan, A.J. Phillips, Madeline Rose Hays, Alexandra Kling, Eric T. Zhao, Sasidhar Madugula, Ramandeep S. Vilkhu, Praful Krishna Vasireddy, Andreas Hierlemann, Guosong Hong, E.J. Chichilnisky, Nicholas A. Melosh

## Abstract

Silicon-based planar microelectronics is a powerful tool for scalably recording and modulating neural activity at high spatiotemporal resolution, but it remains challenging to target neural structures in three dimensions (3D). We present a method for directly fabricating 3D arrays of tissue-penetrating microelectrodes onto silicon microelectronics. Leveraging a high-resolution 3D printing technology based on 2-photon polymerization and scalable microfabrication processes, we fabricated arrays of 6,600 microelectrodes 10-130 µm tall and at 35-μm pitch onto a planar silicon-based microelectrode array. The process enables customizable electrode shape, height and positioning for precise targeting of neuron populations distributed in 3D. As a proof of concept, we addressed the challenge of specifically targeting retinal ganglion cell (RGC) somas when interfacing with the retina. The array was customized for insertion into the retina and recording from somas while avoiding the axon layer. We verified locations of the microelectrodes with confocal microscopy and recorded high-resolution spontaneous RGC activity at cellular resolution. This revealed strong somatic and dendritic components with little axon contribution, unlike recordings with planar microelectrode arrays. The technology could be a versatile solution for interfacing silicon microelectronics with neural structures and modulating neural activity at large scale with single-cell resolution.

## Introduction

With rapid developments in neuroelectronics and optogenetics, it is now possible to record and/or stimulate electrical activity in hundreds of neurons simultaneously^1–3^. This enables investigations into the mechanisms behind neural functions such as motor control and decision making, as well as development of next-generation neural prosthetics with improved performance (*e*.*g*. speech prostheses enabled by decoding high-density neural activity in the sensorimotor cortex)^4–7^. Neuroelectronics is advantageous for developing such technologies compared to optogenetics, because it does not require genetic modification, it relies on simpler setups for signal readout and write-in, and can be applied to most parts of the nervous system^8^. More importantly, thanks to advanced silicon processing, planar arrays with thousands of microelectrodes packed at a spatial density close to that of neurons have recently been demonstrated^9–11^, a significant leap from previous-generation technologies with a little over one hundred channels and coarsely spaced microelectrodes.

However, neurons are typically organized in three-dimensional (3D) structures that are difficult to address with planar silicon microelectronic arrays. Recent innovations to bridge this geometric mismatch include fabricating complementary metal oxide semiconductor (CMOS) circuits into shanks that can be inserted into neural tissues^10,12^, connecting planar electronic arrays to insertable microelectrodes^13–18^, growing silicon or metal pillars as microelectrodes onto CMOS arrays^19–21^, and patterning polymer or oxide pillars that are then metalized to become microelectrodes^22–24^. These technologies leverage the spatial resolution and scalability of microfabrication processes, but unfortunately do not allow a high degree of flexibility or customizability for interfacing to diverse neural populations.

Retinal implants for vision restoration are a key example of a neural interface in which the spatial location of recording/stimulating microelectrodes needs to match the 3D organization of the target neurons. In normal vision, the retinal circuitry extracts information about the visual scene and encodes it into the spatiotemporal patterns of the retinal ganglion cells (RGC) activity known as the RGC “neural code”^25,26^. High-acuity artificial vision thus requires understanding and reproducing this neural code through electrical recording and stimulation at high density. This requires precise placement of the microelectrodes into the RGC layer to avoid unwanted activation of axon bundles^27,28^. Additionally, the contour of the microelectrode array should conform to the natural curvature of the retina, and would ideally be customizable to the individual retina to be implanted. Finally, the electrodes should be able to target the multiple layers of RGCs present around the fovea, requiring the microelectrodes to be of different heights at different regions. The need for high-density recording sites over a two-dimensional area makes this a good match for using planar silicon-based microelectronics, yet a multi-layered microelectrode array is challenging to fabricate using traditional silicon processing schemes.

With its flexible ability to create customizable structures, additive manufacturing (also known as 3D printing) has recently been explored to fabricate neural interfacing microelectrodes^29–32^. However, these technologies require high processing temperatures and have coarse spatial resolution, limiting their ability to fabricate high-density microelectrode arrays and applicability for silicon microelectronics. The recently developed 2-photon polymerization (2PP) forms structures by crosslinking photosensitive polymers through simultaneous 2-photon absorption^33^. This allows 3D printing with sub-micron resolution, a two-order-of-magnitude improvement from current single-photon technologies such as stereolithographic apparatus (SLA), making it an ideal method for directly printing high-density, individually customizable structures as templates for microelectrodes. For example, Brown et al. recently used 2PP to fabricate high-resolution microelectrodes onto the ends of electrical traces on flexible polymer substrates^34^. These flexible devices can be implanted to the surface of the brain, and efforts to record neural activity are currently underway, but the devices are restricted in spatial density (90-µm pitch) and channel count (16) due to the need for contacting each electrode with an electrical trace on the substrate and laser micro-machining performed on each probe to remove the passivation material.

Here, we developed a method to fabricate fully customizable and high-density 3D arrays with 6,600 microelectrodes at 35-μm pitch directly onto advanced silicon microelectronics. This was achieved by combining the customizability and high spatial resolution of 2PP, which allows precise control on the shape and height of each individual microelectrode, with scalable microfabrication processes that eliminate the need for sequential processing on individual electrodes. This process decouples the design and fabrication of the tissue-interfacing microelectrodes from the underlying electronics, allowing for easy experimentation with microelectrode geometries and customization towards specific neural tissues. As a demonstration, we designed the microelectrode array to address the aforementioned challenge of interfacing with RGCs in the retina. We customize the height of the microelectrode array to target RGC somas while avoiding the axon bundle layer upon insertion into the retina. Successful targeting was verified with confocal microscopy and electrical recording *ex vivo*, demonstrating excellent spatiotemporal neural recordings at high density over a large area. This could be a promising solution for high-acuity artificial vision produced by retinal prostheses—the high density and customizability of the array could allow precise activation of RGCs *en masse* to reproduce the natural RGC neural code^27^. Additionally, we believe the application of this technology can be extended to other parts of the nervous system and may be a useful tool for precisely addressing neural circuits at high spatial resolution, enabling large-scale neural interfacing at singlecell and single-cell-type resolution.

## Results

### Scalable, high-density 3D microelectrode array enabled by 2PP

3D printing with 2PP can fabricate customizable structures with high spatial resolution, making it an ideal method for directly printing high-density, individually customizable microelectrodes at scale on silicon. **Fig. 1 (a)** shows our approach to producing individually insulated microelectrodes with conductive tips and controllable height and shape. The non-conductive core of the electrode is printed with 2PP directly onto the pixels of the silicon microelectronics array, followed by metallization, patterning and passivation steps. The cutaway renderings show the layers of materials of the microelectrode after the fabrication process. The microelectronics array used in this work has been fabricated in CMOS 180-nm technology and features 26,400 microelectrode pixels at 17.5-μm pitch, 1024 low-noise (<10 μV) recording channels and 32 stimulation units^35^.

**Figure 1:**
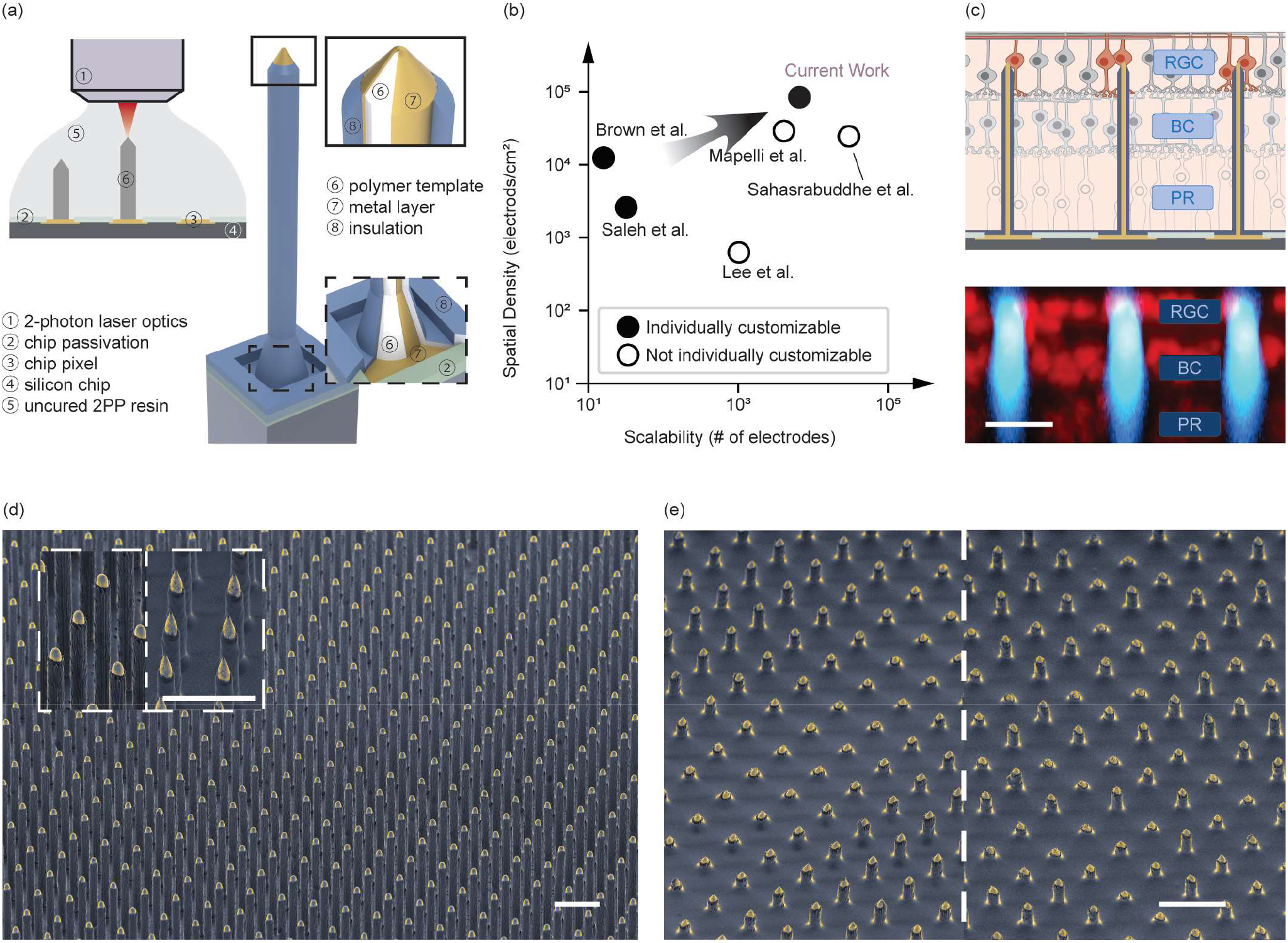
Technology overview. **(a)** 2PP enables direct printing of 3D microelectrodes onto high-density silicon microelectronics (④). (Left) The microelectrode templates (⑥) are first directly written. Through photolithographic patterning a conformal layer of Au (⑦), conformal insulation deposition (⑧), and selective removal of the insulation layer at the microelectrode tips, the templates are converted into individually addressable microelectrodes. (Right) 3D cutaway rendering shows the material layout of the microelectrodes. **(b)** Our technology achieves the highest spatial density and scalability while offering customizability at single-microelectrode level. **(c)** With our technology, we tackle the critical challenge of precisely targeting the RGCs when interfacing with the retina. (Top) We customized the height of the microelectrodes to ensure the microelectrode tip is in proximity to RGC somas. (Bottom) Successful targeting of RGC somas is verified by confocal microscopy. (red – cell nuclei, blue – microelectrodes, BC – bipolar cells, PR – photoreceptors) **(d)** SEM images showing the highly uniform 6,600-microelectrode array that was used to record from the retina. Inset shows an example of customizing tip shapes: the conical shape could be used when easier penetration is prioritized; the hemispherical tips with less exposed area could for more localized recording. **(e)** Our technology allows the height of each microelectrode to be varied to target neurons distributed in 3D. As an example, left shows an array with stripes of microelectrodes with varying heights (10-50 μm); right is an array of randomly distributed microelectrodes with varying heights. These arrays could be used to target RGCs around the fovea that are stacked in 3D (Images in **(d)** and **(e)** were obtained with the samples tilted at 45°; scale bars: 20 μm in **(c)**, 50 μm in **(d)** and **(e)**).

The fabrication process consists of the following steps (see Materials and Methods, Table 1 for detailed information about each step, and Supplementary Fig. S1 for illustration of the fabrication process and the scanning electron microscopy (SEM) images of the sample after each step). Following 2PP of the microelectrode template on the pixel array, the device was sputter-coated conformally with 10 nm Ti and 75 nm Au. The metal bilayer was then patterned by a Ge lift-off sacrificial layer defined through standard photolithography prior to the 2PP step. This lift-off process allows each microelectrode to be electrically connected only to the CMOS pixel underneath. The microelectrodeCMOS assembly was then passivated by Parylene-C, and the tips were exposed at scale by oxygen plasma etching. This process was highly uniform, showing less than 5% variation in the area of the exposed tips. Despite the small size and high aspect ratio of the 2PP structures, they withstood the mechanical and thermal stress along the fabrication process, allowing highly uniform arrays to be fabricated.

**Table 1.**
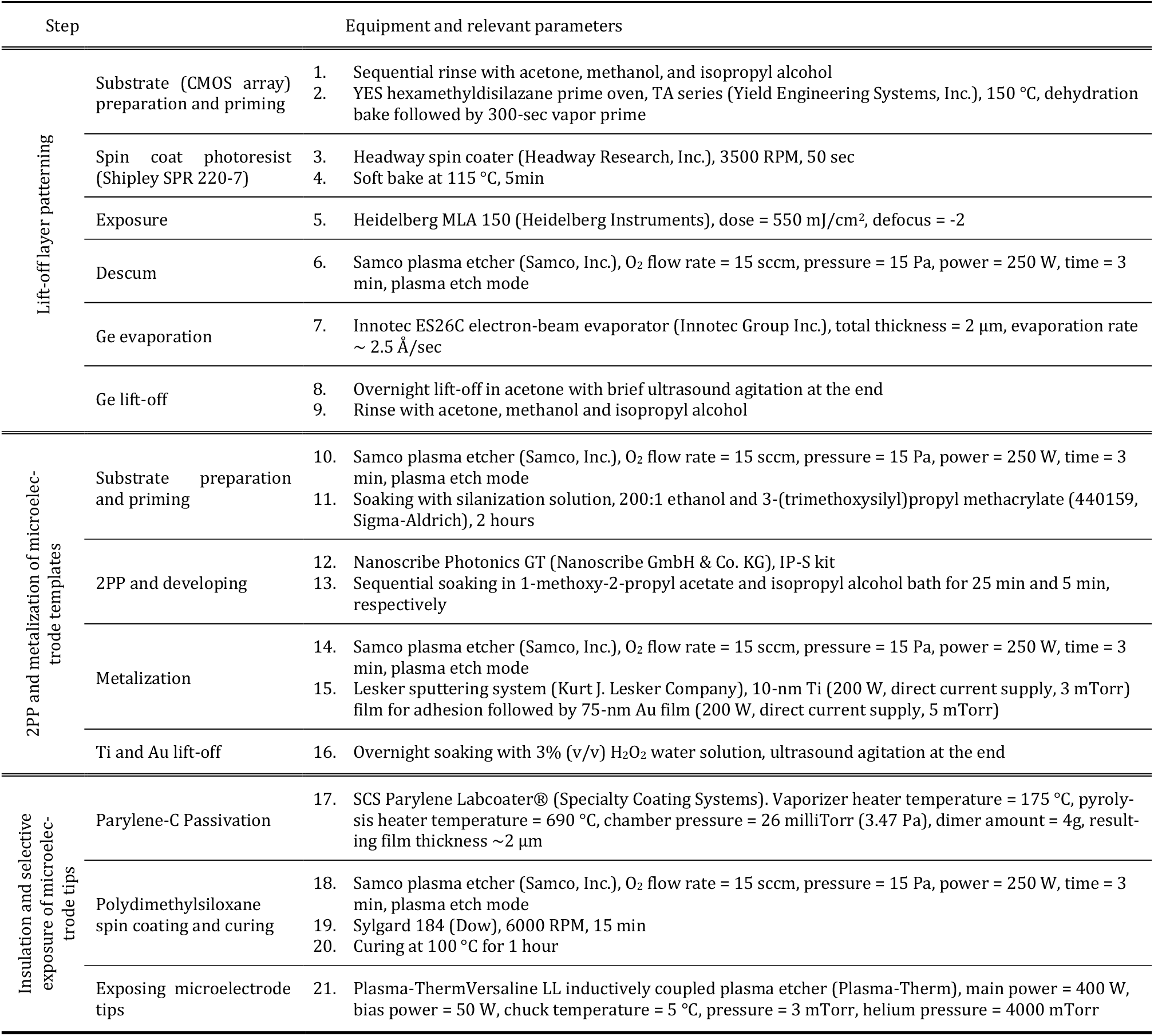
Fabrication Process of the Direct-Print Microelectrodes onto CMOS Arrays

This combined high-resolution 3D printing and silicon microfabrication processes achieved the highest scalability and spatial density for customizable microelectrodes compared to existing technologies (**Fig. 1 (b)**), as well as the capability to customize the array to target RGC somas while avoiding the axon bundle layer (**Fig. 1 (c)**). **Fig. 1 (d)** shows the fabricated high-density (35-μm pitch, equivalating 84,100 microelectrodes per cm^2^) and high-aspect-ratio (110-μm height, 10-μm diameter) arrays consisting of 6,600 microelectrodes. We demonstrate the capability to precisely control the geometry of the microelectrodes with two examples: **Fig. 1** (**d**, inset) shows that the shape of the electrode tips can be optimized to tailor different requirements – for example conical tips for easier tissue penetration and hemispherical tips for more localized sensing; **Fig. 1 (e)** shows arrays of microelectrodes with varying height, which, unlike planar arrays or protruding electrode arrays with uniform height, can be used to bypass certain structures and address neurons distributed in 3D.

### Microelectrode characterization for neural recording

We characterized and modified the electrode-liquid interfacial impedance of the microelectrodes to ensure they match the impedance requirement of the CMOS circuitry^35^. We first measured the impedance with electrochemical impedance spectroscopy (EIS). As EIS cannot be directly performed on individual electrodes on the CMOS array due to limitations of the array that we used, we separately fabricated the microelectrodes on a 16-channel planar microelectrode array (16MEA, shown in **Supplementary Fig. S2**) using the same processes described above. The microelectrodes showed high amplitude of impedance at 1 kHz (|*Z*|_*1k*_ = 3.8 MOhm ± 1.5 MOhm) due to the small surface area of the exposed tips and the relatively high electrochemical impedance of gold. To lower the impedance, we electroplated platinum-black (Pt-black) onto the tips of the microelectrodes, which lowered |*Z*|_*1k*_ by 2 orders of magnitude (|*Z*|_*1k*_ = 23 kOhm ± 6.7 kOhm). **Fig. 2 (a)** shows the distribution of |*Z*|_*1k*_ before and after electroplating Pt-black. Supplementary **Fig. S2** and **S3** shows the set-up of EIS measurement and Pt-black electroplating, bode plots between 1-100kHz before and after Pt-black electroplating, as well as SEM image of the electrode tip coated with Pt-black.

**Figure 2.**
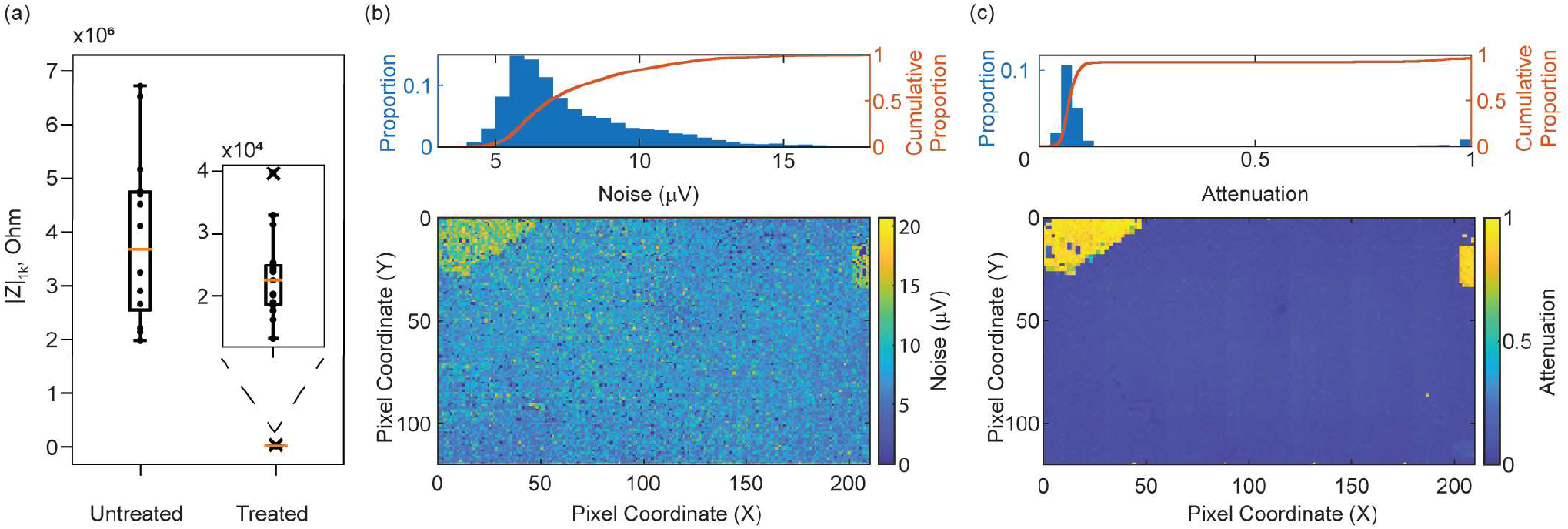
Suitability of the microelectrode array for neural recording. **(a)** The microelectrode tips can be modified post-fabrication (*e*.*g*. electroplated with Pt-black) to lower the electrode-liquid interfacial impedance. Box plot (orange line — median; box limit — first to third quartile; whiskers — box limit ± 1.5x interquartile range; dots — individual data points; crosses — outliers) shows the distribution of |Z|_1k_ before and Pt-black electroplating. **(b)** Noise and **(c)** signal attenuation histogram and map of the microelectrodes fabricated on the CMOS array. Noise and signal attenuation are highly uniform across the entire array, with anomaly at the top left and right corners caused by accidental damages to the microelectrodes during handling, as can be seen from **Supplementary Fig. S4** showing the packaged array.

After developing this modification process, we verified that it can be used on the CMOS array fabricated with the microelectrodes, and the CMOS array remains fully functional after the fabrication and modification processes. The pixels on the CMOS array were connected to an external current source to apply the Pt-black electroplating current to the microelectrode tips. Uniform electroplating was confirmed by noise and signal attenuation measurements in phosphate buffered saline (PBS, 1X). The configuration for electroplating and optical microscopy images of the electrode tips coated with Pt-black are available in **Supplementary Fig. S2** and **S3**. Noise after electroplating was calculated as the root mean square (RMS) value of a blank recording band-passed between 300 and 4,000 Hz. **Fig. 2 (b)** shows the noise distribution of one array with 85.2% of the microelectrodes having noise below 10 µV. Signal attenuation was measured by injecting a 5-mV sinusoidal signal at 1 kHz into the PBS through a Pt electrode connected to a function generator and measuring the amplitude of the recorded signal. **Fig. 2 (c)** shows the minimal and highly uniform signal attenuation across the entire array. These results deem the microelectrode array suitable for neural recording.

### Penetration into the retina and high-density recording from RGC somas

High-density electrodes often pose challenges for penetration into neural tissues because of the “bed-of-nails’’ effect that distributes the insertion pressure over the entire array and causes tissue dimpling and damage^36^. Here, with the small microelectrode tip diameters (<5 µm) and low volume of tissue displacement — the volume fraction of the microelectrodes is 1.6% — we were able to penetrate retina tissue with our microelectrode array as verified by confocal microscopy. A segment of isolated macaque retina approximately 2 mm in diameter was first positioned onto the microelectrode array with the photoreceptor side facing the array, and a custom-built pressing apparatus was used to apply uniform pressure to insert the microelectrodes into the retina (**Supplementary Fig. S5**). After visually confirming that the retina was uniformly flattened and the microelectrodes became visible, we removed the pressing apparatus and dyed the nuclei of the retinal neurons with DRAQ5™. The shorter-wavelength auto-fluorescence from the passivation layer of the microelectrodes allowed them to be distinguished from the cell nuclei, and Z-stacks were obtained with confocal imaging.

The imaging results revealed that the microelectrode array through most of the retina, with the microelectrode tips precisely positioned in the RGC layer and the array causing minimal disturbance to the surrounding cells. **Fig. 3 (a)** is a 3D rendering from a z-stack of confocal imaging showing the retina penetration (a 3D animation is also available in the **Supplementary Video**). The orthogonal view from the same z-stack is shown in **Fig. 3 (b)**. The *xz* view confirms that the microelectrodes penetrated the retina with the tips positioned at the RGC layer. The three main layers of retinal neurons – photoreceptors, bipolar cells, and retinal ganglion cells (RGCs) – are clearly visible on both the *xz* and *yz* views. To quantify the position of the microelectrode tips relative to RGCs, we measured the positions of RGCs in the z-stack. **Fig. 3** (**c**, top) shows the microelectrode tips were positioned near the region where RGC density peaks. Importantly, after calculating RGC density (number per mm^3^, binned every 10 μm by distance to selected microelectrode tips or RGCs in between), we found that the high density of the microelectrode array did not significantly disturb the RGC distribution as shown by the negligible variation in RGC density. This finding is consistent with the low volume displacement (1.6%) of the penetrating electrodes. It is also worth noting that the array remains largely intact after tissue penetration as can be seen in **Fig. 3 (a)** and **(b)**, as well as in an optical microscopy image of the array obtained after the tissue had been removed from the array (**Supplementary Fig. S5**). This demonstrates the mechanical robustness of the microelectrode array.

**Figure 3.**
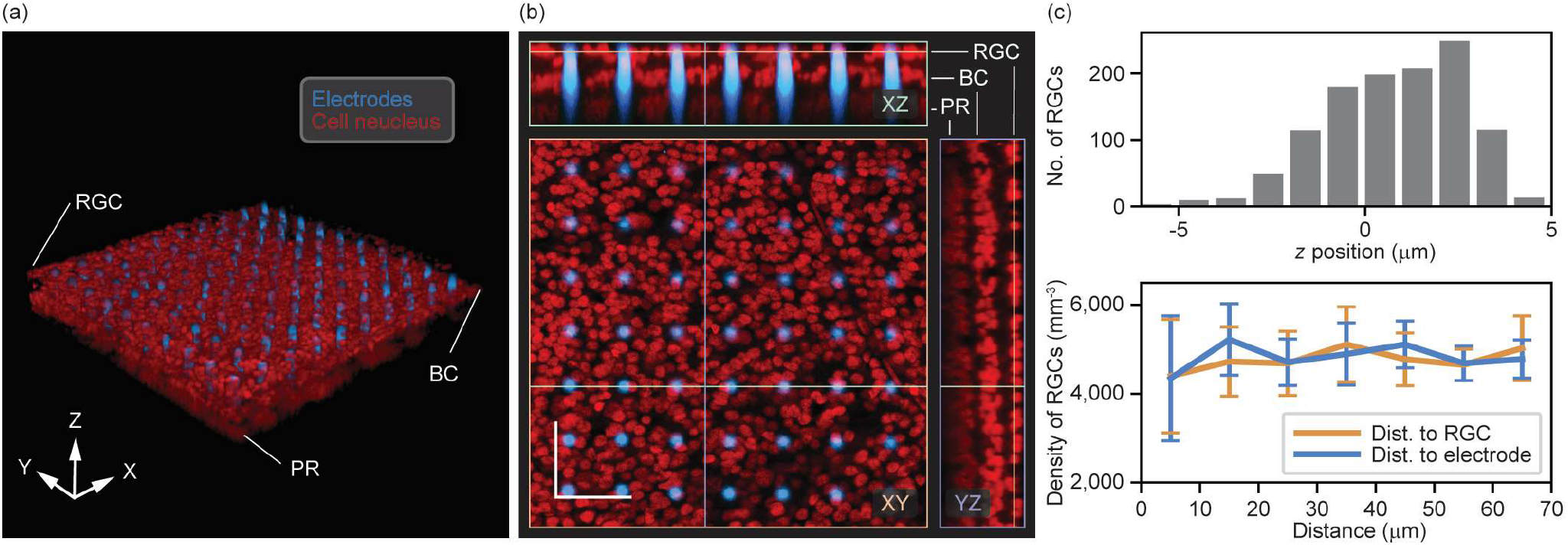
Tissue penetration with minimal disturbance. **(a)** 3D rendering from a z-stack obtained with confocal imaging. The autofluorescence of the passivation layer of the microelectrodes allow them to be distinguished from the cell nucleus that were dyed with DRAQ5™. **(b)** Orthogonal view of the z-stack with the *XY* view showing the high-density microelectrode array surrounded by RGCs. The *XZ* view verifies tissue penetration and the location of the microelectrode tips being in the RGC layer. The layered structure of retinal neurons is visible in both *XZ* and *YZ* views. Positions of the orthogonal planes are indicated by the color-coded lines. Scale bar: 50 μm. **(c)** (Top) vertical distribution of RGC relative to the microelectrode tips (*z* = 0) shows that the microelectrode tips are positioned near RGC peak density. (Bottom) Density of RGCs as a function of distance to selected reference points (*i*.*e*. microelectrodes and RGC in between, error bars: standard deviation of multiple measurements across the image). There is no significant variation in RGC density as distance increases or between the two measurements with different reference points. This shows that the microelectrode array minimally disturbs RGC distribution.

To test the suitability of the electrode array for electrophysiological experiments, we obtained recordings from a 2×2 mm segment of isolated rat retina. We inserted the microelectrode array from the photoreceptor side of the retina by applying force while monitoring the recorded electrical activity. Because the photoreceptor cells and most bipolar cells of the retina do not generate action potentials, we inferred that the probe tips were near the RGCs when action potentials were observed. We then recorded spontaneous RGC activity using subsets (∼900) of the 6,600-microelectrode array at a time. Example bandpass filtered spikes recorded from the electrodes are shown in **Fig. 4 (a)**, with somatic spike amplitudes in the 66.2 ± 32.0 µV range, and typical baseline recording noise (combined instrument and physiological noise) on each electrode of roughly 19.8 ± 1.3 µV RMS. The data were spike sorted using Kilosort 2^37^, and putative RGCs showing physiologically realistic refractory period and firing rate (*n* = 34) were analyzed further.

**Figure 4.**
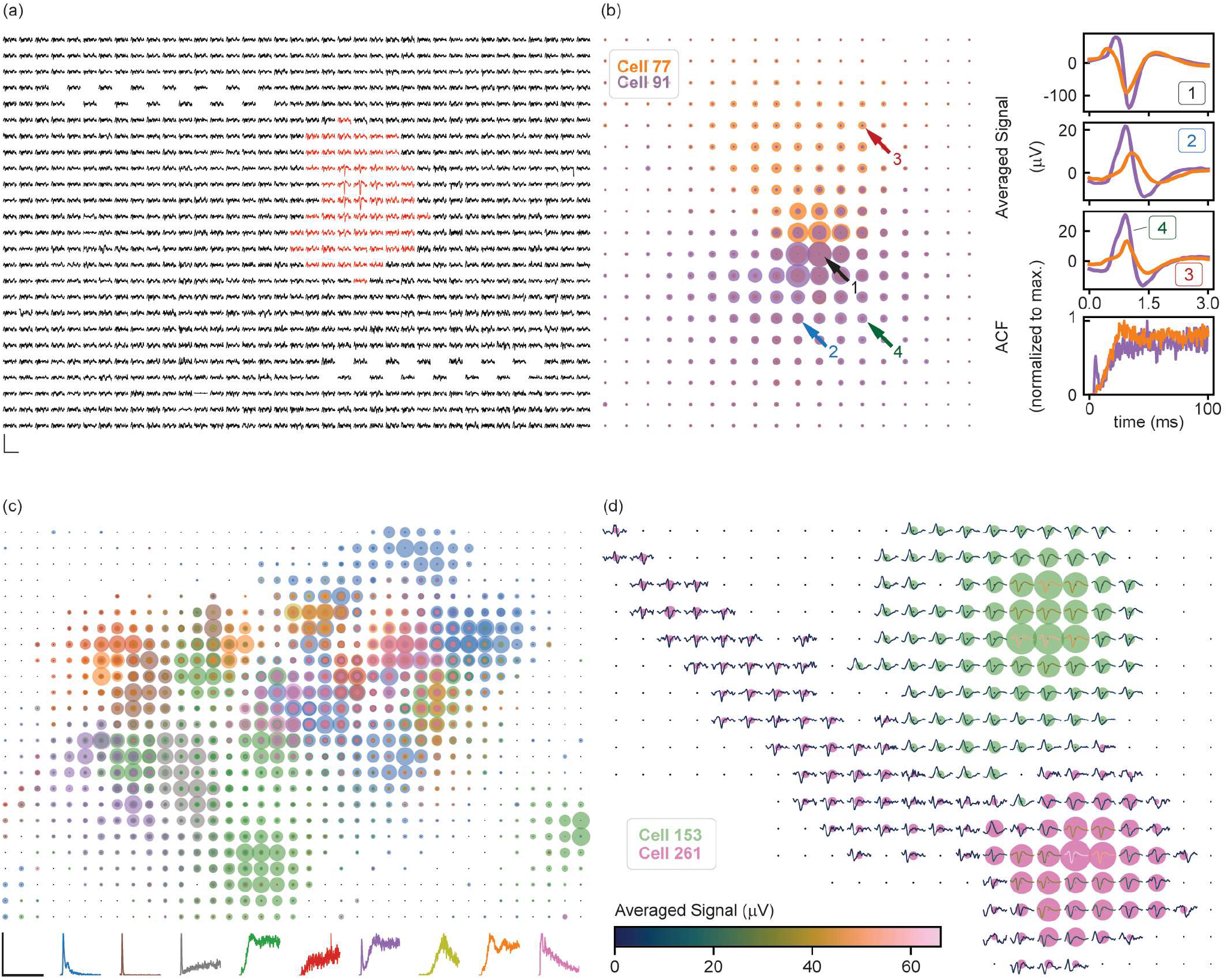
High-density and large-scale recording from rat RGCs ex-vivo. **(a)** Traces of raw signals recorded with 897 microelectrodes, around the firing time of an RGC located near the center of the sampled microelectrodes. Red traces are recorded with microelectrodes to which the EI from the firing RGC have been mapped. Several sites show no recordings near the top left and bottom right locations due to routing limitations of the CMOS array. Scale bar: horizontal – 5 ms, vertical – 100 μV. **(b)** EIs of two example cells. Each dot on the EI is centered around the microelectrode position. The sizes of the dots correspond to the amplitude of the averaged voltage deflections recorded around the spiking times of a cell. The highdensity recording allows RGC in proximity to be clearly distinguished. Cell 77 and Cell 91 have overlapping somatic activity, but different dendritic field spatial extents, as shown by their EIs and averaged voltage deflections recorded from locations indicated by the arrows. Their ACFs suggest distinct temporal activity patterns. **(c)** Overlaid EIs of all the analyzed RGCs. The ACFs of the RGCs indicate distinct groups of temporal firing patterns, with representative ACF (normalized to respective maximum) from each group shown at the bottom. The EIs are color-coded according to the ACFs. ACF scale bar: horizontal – 120 ms, vertical – 1 (unitless). **(d)** To show the difference among somatic, dendritic, and axonal signals, the EIs from two example cells are overlaid with the averaged voltage deflections across 3 ms and normalized and color coded by their amplitude. Most of the recorded RGCs show dominantly somatic and dendritic activity, such as cell 153. Of the cells that do show axonal signals (*e*.*g*. cell 261, evidenced by the axon trajectory on the EI), the amplitude is much smaller compared to those recorded with planar electrodes.

The high spatial density and scale of this recording allowed us to obtain a high-fidelity representation of the electrical signals in each identified RGC. Using a previously reported method^38,39^, we calculated the electrical image (EI) of each RGC (the average spatiotemporal voltage signal recorded across the array during a spike) to identify the soma, dendrite and axon signals recorded across the array. The autocorrelation function (ACF) of the RGC spike train was also calculated to characterize the temporal pattern of the spontaneous activity. As an example, **Fig. 4 (b)** shows the EIs of cell 77 and cell 91, which can be clearly distinguished despite their proximity because the somatic and dendritic signals exhibit different waveforms and amplitudes, and because the two cells exhibit different dendritic field spatial extents. Additionally, the ACFs reveal different temporal patterns of activity in the two cells. The EIs of all the analyzed RGCs are plotted in **Fig. 4 (c)**. Notably, the RGCs in **Fig. 4 (c)** can be grouped according to the shapes of their ACFs, potentially revealing distinct RGC types^40,41^. The representative ACF in each group is shown in **Fig. 4 (c)**, and the EIs are color-coded according to their corresponding ACF. These results demonstrate that the penetrating microelectrode array can obtain high-quality recordings with high temporal-spatial resolution.

The EIs observed with the penetrating microelectrode array differed qualitatively from those observed using flat electrode arrays. While strong dendritic and somatic signals (identified by their unique waveforms (**Fig. 4 (d)**) were observed, only 7 of the 34 identified cells showed axonal signals as indicated by the presence of a long propagating projection from the somatic compartment of the EI (**Fig 4 (c)**). We attribute this to the fact that the penetrating electrode tips are closer to the RGC somas and further away from the axon bundle layer compared with planar arrays. This finding implies that cross interference with the axon bundle layer can be significantly reduced by inserting the microelectrodes specifically into the RGC soma layer. This has not been demonstrated by existing technologies, and is enabled by our capability to fabricate high-density 3D microelectrode arrays that can penetrate tens of µm into neural tissue and can be customized to address the spatial distributions of neurons in 3D.

## Discussion

We demonstrated that 2PP combined with standard microfabrication processes is a versatile method for directly fabricating high-density penetrating microelectrodes onto silicon microelectronics at scale. The method offers a high degree of customizability, which allows us to address the challenge of targeting RGC somas while avoiding the axon layer. We verified successful retina penetration and avoidance of the axon layer with fluorescent microscopy and electrical recording. The high-quality recording with our array allows precise mapping of retinal neurons, with a spatial resolution revealing activities from various cellular compartments within a single cell.

2PP has shown limited application in neuroelectronics because most of the materials available for 2PP are mostly non-conductive polymers. Although 2PP of conducting polymers is possible, the trade-off between conductivity and structural deformation during printing limits their potential for fabricating microelectrodes with high precision^42^. Our process proves to be compatible with silicon microelectronics that are often sensitive to high temperatures and electric discharging events. The electrochemical properties of the microelectrodes were verified to be highly uniform and suitable for large-scale neural recording. Although electroplating of platinum black was required to lower the electrode-saline interfacial impedance, it is possible to directly deposit low-impedance materials, such as titanium nitride or iridium oxide, during the sputter-coating and lift-off steps of the process. This feature makes our process completely compatible with any integrated circuits including those that do not support on-chip electroplating.

To the best of our knowledge, tissue penetration with 3D electrode arrays of this density has not been reported previously. We successfully inserted these high-density microelectrodes into retinal tissue while causing minimal displacement of RGCs, resulting in high-quality recordings. The shape of the microelectrode tips could be optimized for insertion in other applications^34^. This is a direct benefit from the versatility of 2PP that is impossible to reach with conventional microfabrication processes.

Tissue penetration proved beneficial for RGC recordings, presumably because penetrating microelectrode tips are closer to the neurons than planar electrodes positioned on the surface of neural tissues^43^. Furthermore, the recordings demonstrated that penetrating electrodes can reduce interference from axons. For future retinal implant efforts aimed at vision restoration, these penetrating electrodes could also be used to selectively electrically stimulate the axonal initial segment near a RGC soma and avoid inadvertently stimulating passing axons from untargeted RGCs.

Combined with previous work that allows identification of RGC types based on their spontaneous activity^41, 44^, it may then be possible to electrically stimulate with cell-type specificity, allowing accurate reproduction of the natural RGC neural code^45^.

The ability to penetrate through several layers of neurons and neuropil with high-density electrodes may prove important for interfacing with the fovea – a region of the retina consisting of RGCs densely packed in several layers and responsible for central high-acuity vision. The flexible fabrication method can be used to address the distinct layers with electrodes of varying length, while also accommodating the complex curved surface of the foveal region. This could enable restoration of central vision, which is critical for most visual tasks.

Together these findings indicate that, in the retina, the penetrating microelectrodes provide a novel way to record (and potentially stimulate) more selectively than was previously possible, and in the future may be important for accessing other multi-layer neural ensembles. These advantages for neural interfacing may be advantageous for targeting other brain structures such as the cortex where neurons are organized by layers. Additionally, we believe multimodal neural interfacing is possible with our technology. For example, 2PP can be used to fabricate microfluidic channels together with the micropillars. This would enable high-density and 3-dimensional neural interfacing using both electrical and chemical signals, which could be an emerging technology for various applications including such as drug screening and studying neural or disease development.

## Acknowledgements

We thank Martin Breidenbach, Stuart Cogan, Nofar Hemed, Swaroop Kommera, Phil Himmer, Bui Cat-Vu Huu, Seungbin Jeong for their valuable inputs on developing the microfabrication processes, Ryan Samarakoon and Erin Moon for supporting retina dissection and recording experiments, and Gordon Wang for discussions and guidance on confocal imaging of the retina. This work was funded by the Wu Tsai Neuroscience Institute, Stanford University (Big Ideas grant, to N.A.M. and E.J.C.), and by the National Eye Institute (grant No. 5R01EY032900, to E.J.C. and N.A.M.). This work was performed in part in the nano@Stanford labs, which are supported by the National Science Foundation as part of the National Nanotechnology Coordinated Infrastructure under award ECCS-2026822, and in Stanford Neuroscience Microscopy Service. This paper was typeset with the bioRxiv word template by Ebbesen.

## Author contributions

P.W., N.A.M., and E.J.C. conceived the overall study. P.W., N.A.M., and G.H. designed, and P.W. performed experiments on fabricating the microelectrode array with additional advice from E.T.Z., H.U., and A.H. P.W., E.G.W., and E.T.Z. characterized the microelectrode array with advice from H.U. E.J.C. E.G.W., A.J.P., M.R.H., P.W., A.K., S.M., R.V., and P.K.V. designed, and E.G.W., A.J.P., M.R.H., P.W., A.K., S.M., R.V., and P.K.V. performed experiments and analyzed data on retina penetration, confocal microscopy, and recording. P.W., N.A.M., E.J.C., and E.G.W. wrote the manuscript with input from all other co-authors.

## Materials and Methods

### Fabrication, packaging, and SEM imaging of the microelectrode array

The CMOS array contains 25,400 pixels at 17.5-μm pitch. The Ge lift-off layer was defined such that four pixels are used to address one microelectrode, allowing 6,600 microelectrodes to be fabricated per array. 2PP directly on the pixel was challenging due to high reflectance of the metal surface, so each microelectrode template was printed on the passivation next to the pixel. The metalization step established electrical connection between the microelectrode and the pixel. Table 1 summarizes the process and relevant parameters for fabricating the microelectrode array.

The CMOS arrays fabricated with the microelectrodes were wirebonded to custom printed-circuit boards (PCBs) and packaged as described previously^35^. Briefly, after wire-bonding, a ceramic chamber ring was first positioned around the wire-bonds and glued to the PCB using a two-part epoxy (EPO-TEK^®^ 353ND-T, Epoxy Technology, Inc.). Then, the same epoxy was applied around the microelectrode array in the form of a 1-mm line dispensed through a syringe. The epoxy was partially cured at 80 °C for 1 hour. After the assembly cools down from the baking, a different epoxy with thinner consistency (EPO-TEK^®^ 353ND, Epoxy Technology, Inc.) was poured into the space between the previously drawn epoxy line and the ceramic ring to cover the exposed wire-bonds, with the line drawn around the microelectrode array preventing the uncured epoxy from flowing onto the array. Finally, the epoxy was cured with a temperature-profiled baking procedure, where the temperature was first held at 80 °C for 30 minutes, and then ramped to 150 °C within 30 minutes and held for 1 hour. The assembly was slowly cooled over a few hours and ready for testing.

SEM imaging of the fabricated arrays was conducted with a field-emission scanning electron microscope (Quanta 250 FEG, FEI company) using acceleration voltage = 2 kV, spot size = 3, and working distance ∼ 10 mm. The images were false colored using gradient mapping in Adobe Photoshop 2023.

### Characterization and modification of the microelectrode array for neural recording

The CMOS array used in this work does not allow direct ohmic contact to the fabricated microelectrodes to perform EIS, so we separately fabricated the microelectrodes on the custom-designed 16MEA using the same process described above. The microelectrodes are electrically connected to the contact pads on the periphery of the 16MEA.

EIS was measured with two-electrode configuration using a potentiostat (SP-200, BioLogic). A droplet of PBS (1X) was positioned to submerge the microelectrode array, and a Pt electrode (327492, Sigma-Aldrich) was submerged in the same droplet and connected to the counter and reference electrode port on the potentiostat. The contact pad on the 16MEA was connected to the working electrode port and to inject sinusoidal signals (amplitude = 10 mV, injected at 10 evenly spaced frequencies across each decade, total frequency range 1 Hz-100 kHz). The full-spectrum EIS across 16 microelectrodes on the 16MEA was measured and computed using EC-Lab® (BioLogic). One microelectrode was damaged during Pt-black electroplating and was not measured post-electroplating.

To lower the electrode-electrolyte interfacial impedance, Pt-black electroplating on the 16MEA was conducted with the same configuration. Instead of PBS (1X), a droplet of Pt-black electroplating solution containing 3 wt% chloroplatinic acid hexahydrate (206083, Sigma-Aldrich) and 0.3 wt% lead(II) acetate trihydrate (215902, Sigma-Aldrich) was used. Electroplating was performed with a constant voltage (−1.1 V against the Pt wire counter electrode) applied to one microelectrode each time (duration was controlled by the total charge transferred *Q*_*TOT*_< 800 nC), and the 16 microelectrodes were electroplated sequentially. After electroplating, the array was rinsed with deionized (DI) water.

Pt-black electroplating on the packaged CMOS array followed the same procedure as previously reported^35^. In short, after filling the chamber with the Pt-black electroplating solution, a Pt wire connected to an external current source (2410, Keithley) was submerged in the solution. Utilizing the stimulation circuity on the CMOS array, all pixels were connected to a common current sink in order to flow the electroplating current. A constant current of -0.6 mA was applied through the Pt wire into the solution for 1 min. Finally, the solution was removed and the chamber was rinsed with DI water.

Noise and signal attenuation of the microelectrodes were measured using the same method described previously^18^. For noise measurement, with the chamber filled with 1X PBS, a blank recording was obtained with the amplifier gain of the CMOS array set to 512, and noise was calculated by the root mean square amplitude of the blank recording. Attenuation measurement was conducted by injecting a 5-mV, 1-kHz sinusoidal signal into the 1X PBS and recording with the amplifier gain set to 28. Attenuation was quantified using the relative amplitude of the recorded versus the injected signal.

### Tissue preparation, penetration test, and confocal microscopy

Isolated rhesus macaque monkey (*Macaca mulatta*) retinas were used for the tissue penetration test. Tissue preparation followed previously reported protocols^38^ and was in accordance with institutional and national guidelines. Briefly, eyes were removed from terminally anesthetized macaque monkeys, immediately followed by the removal of the anterior portion of the eyes and vitreous. Tissues with size of 2 mm by 2 mm were then dissected. Perfusion with Ames’ solution (A1420, Sigma-Aldrich, 31–36 °C, pH = 7.4, bubbled with 95% O2 and 5% CO2) continued throughout the preparation.

To test and verify retinal tissue penetration, identical dummy arrays were fabricated on bare silicon substrates using steps 10-20 described above without the metal lift-off step (step 14). The dummy array was secured to a petri dish that was used as a chamber for tissue penetration testing. The prepared retinal tissue was transferred to the testing chamber filled with Ames’ solution, and positioned onto the microelectrode array with the photoreceptor side facing downward. A custom-built pressing apparatus consisting of a micromanipulator, a support arm, and a cylinder with nylon mesh (9313T48, McMaster-Carr) stretched taut on the end facing the retinal tissue was used to uniformly apply pressure on the retinal tissue. While monitoring through a stereoscope, the cylinder was slowly lowered by the micromanipulator to insert the microelectrodes into the tissue from the photoreceptor side. After visually confirming that the tissue was in contact with the substrate of the dummy array, the pressing setup was carefully removed.

For confocal microscopy, the tissue embedded in the microelectrode array was first fixed with 1X PBS containing 4% paraformaldehyde (8.18715, Sigma-Aldrich), then rinsed with 1X PBS twice, each lasting 20 min, and finally stored in 1X PBS. Then, DRAQ5™ (5 mM, 62251, ThermoFisher) was added to the preparation and gentle agitation was applied for 30 min. Finally, the dummy array together with the retinal tissue was added with a mounting medium (ab104135, Abcam) and prepared into a microscope slide. Confocal microscopy was performed on ZEISS Airyscan2 LSM 980 installed with a Plan-APO 10x objective using a 639-nm excitation laser. The signals from the retinal tissue and the microelectrodes were filtered by the program (ZEN 3.3, Blue Edition) built-in DRAQ5™ and DAPI filters, respectively. Z-stacks with 1-μm spacing between adjacent slices were obtained from regions of interest.

Each slice of the Z-stack was adjusted by the “enhance contrast” process in ImageJ, with the fraction of saturated pixels set to 0.35%. The 3D rendering was generated with the “3D viewer” plugin in ImageJ. The XZ and YZ images of the orthogonal view were denoised with an ImageJ plugin developed by Mannam *et al*^46^.

### x-vivo recording from rat retina and calculation of electrical images

Rat retinal tissue was prepared and the CMOS array with microelectrodes were inserted using the same method described above. To position the microelectrode tips near the RGCs, the microelectrodes were slowly inserted from the photoreceptor side while simultaneously recording, and insertion stopped when putative action potentials obtained with signal thresholding were observed. Subsets (∼900) of the 6,600 microelectrodes were used to record spontaneous RGC activity at a time as the CMOS array allows simultaneous recording with up to 1,024 channels. The CMOS array amplifier gain was set to 1024.

Recording data analysis and electrical images of the RGCs used the method reported by Li *et al*. and Litke *et al*.^38,39^ Briefly, the recorded signals were bandpass-filtered with a finite impulse response filter with a passband between 300 Hz and 6000 Hz, and then spike-sorted with Kilosort 2 to identify candidate neurons and their spiking time. The autocorrelation function and the average firing rate of the neurons were calculated, and only neurons showing refractory period and realistic average firing rates were included in analyzing the electrical images. To compute the electrical images of an identified RGC, we averaged – across all the spike times – the signals recorded by each microelectrode within the time window (−1 ms to 2 ms) around the peak negative potential. The amplitude of the averaged signal is represented by the size of the discs plotted at the coordinate of each microelectrode. This generates a visual mapping showing the RGC’s activity in the microelectrode space, and we repeated the calculation for all identified neurons.

