## Supplementary Materials for "Direct-print three-dimensional electrodes for large-scale, high-density, and customizable neural interfaces"

### Fabrication Process

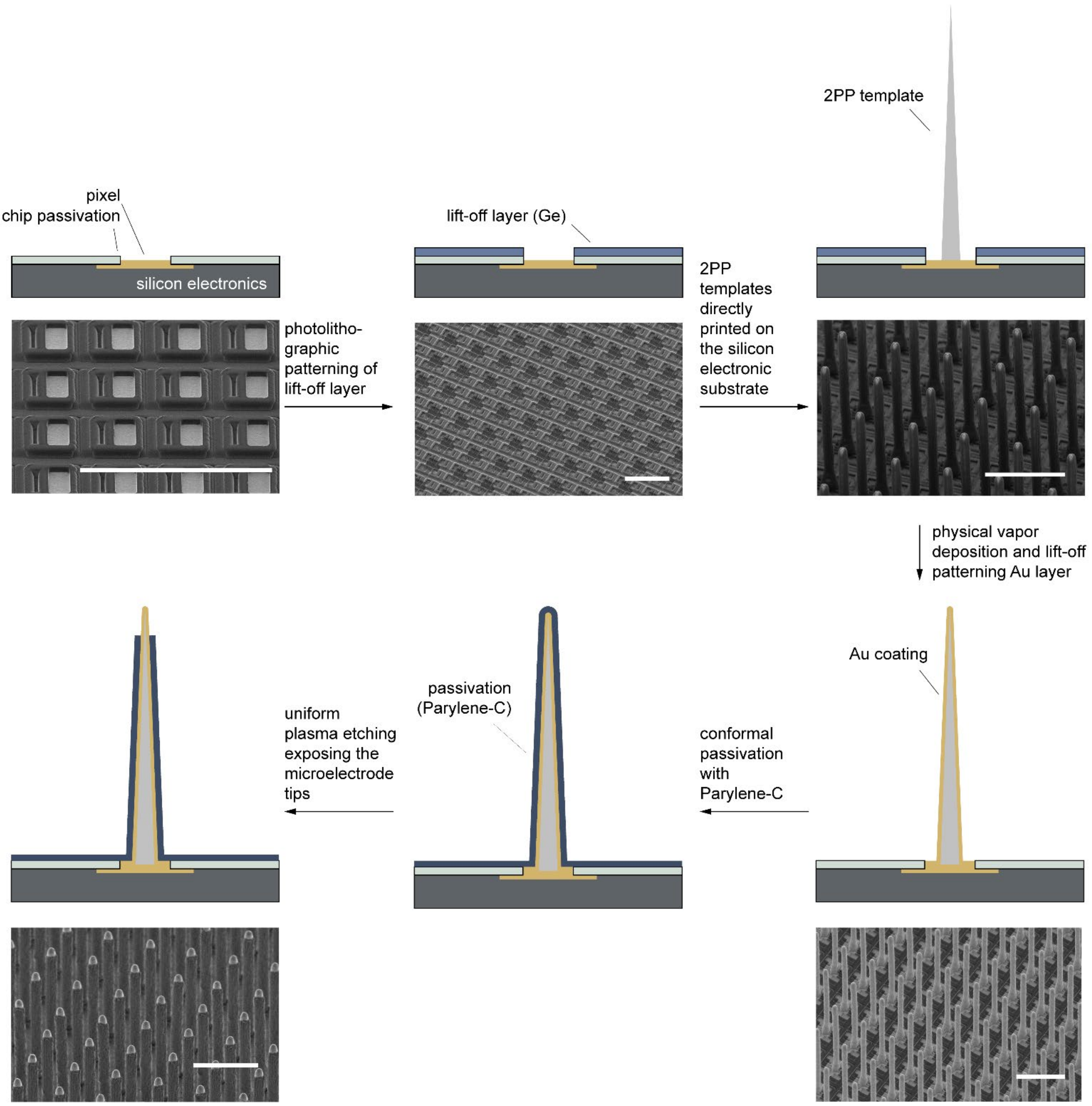

**Fig. S1 Fabrication process and step-by-step SEM images** (scale bars: 50 μm)

### Electrode development for neural recording

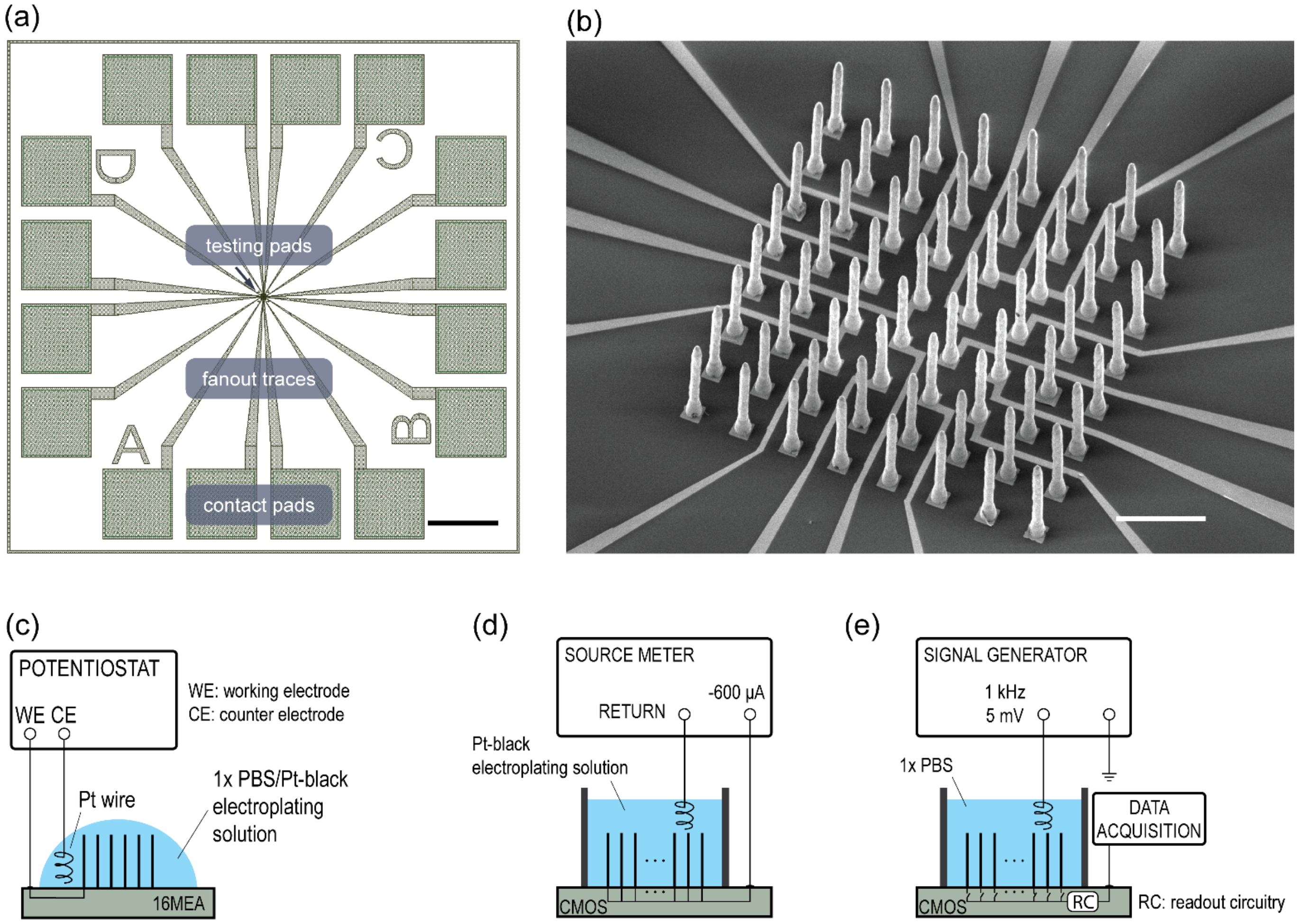

**Fig. S2 16MEA and setups for electrode development**
**(a)** Layout of testing pads and contact pads on the 16MEA. Scale bar: 3 mm. **(b)** SEM image showing an 8 x 8 microelectrode array printed onto the 16MEA, of which 16 are electrically connected by the fanout traces to the contact pads for testing; image was acquired after the Au lift-off step. Scale bar: 50 µm. **(c)** Setup for EIS and Pt-black electroplating using the 16MEA. **(d)** Setup for Pt-black electroplating on the packaged CMOS array printed with the microelectrode array. All microelectrodes are electrically shorted to an external source meter to allow for simultaneous electrodeposition across the entire microelectrode array. **(e)** Setup for signal attenuation measurement using the packaged CMOS array printed with the microelectrode array. Given the limited number of recording channels (1024), the on-chip switching circuits were used to sample tiling subsets of the 6,600 microelectrodes in order to map the attenuation of the entire array. For details of the readout circuitry, please refer to Müller *et al.*, 2015 referenced in the main text.

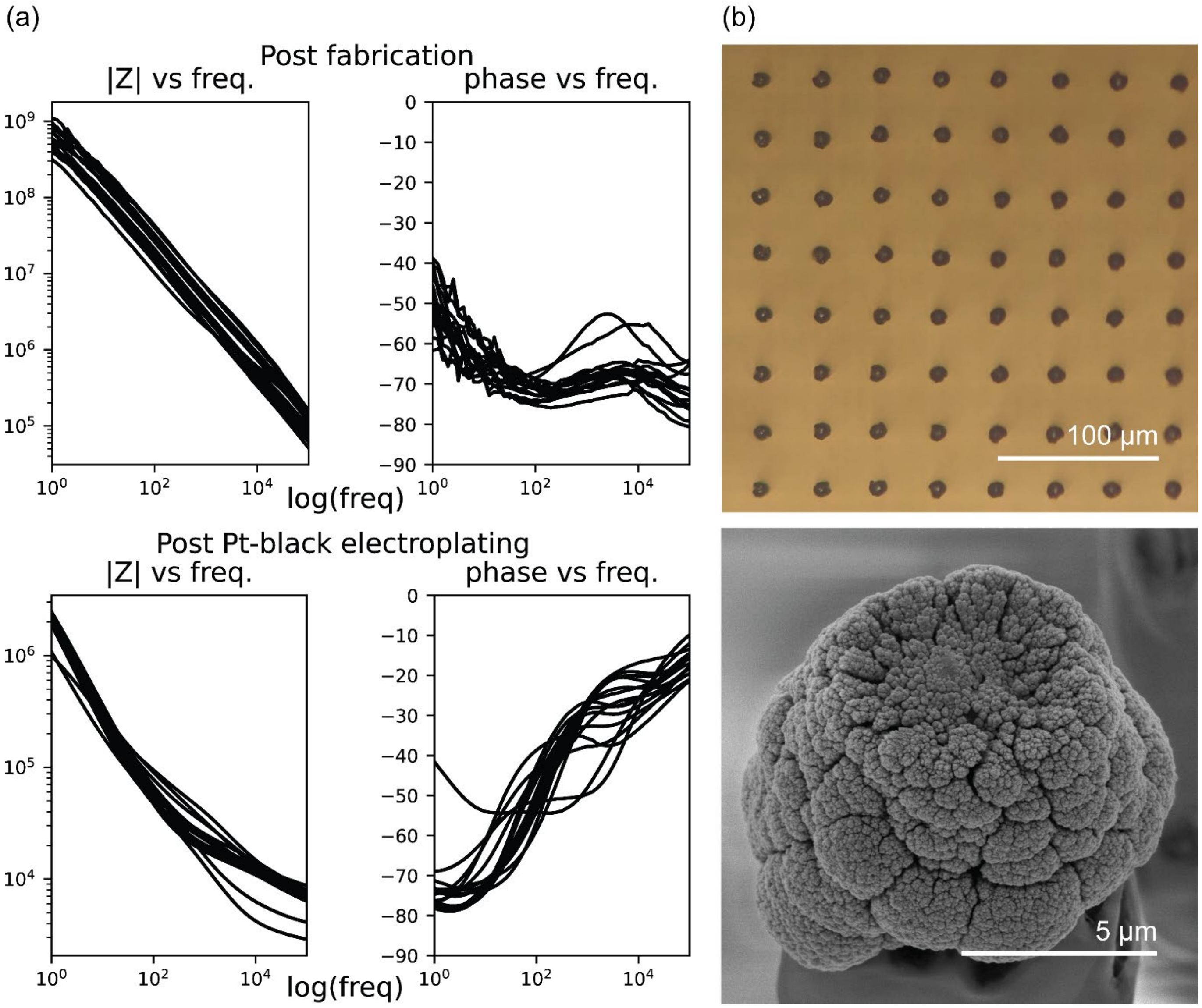

**Fig. S3 Electrode characterization for neural recording**
**(a)** Bode plot (*n* = 16) for a frequency range of 1 Hz-100 kHz, before and after Pt-black deposition; **(b)** Optical micrograph (top) and SEM image (bottom) of microelectrode tips coated with Pt-black

### Packaged device and retina recording setup

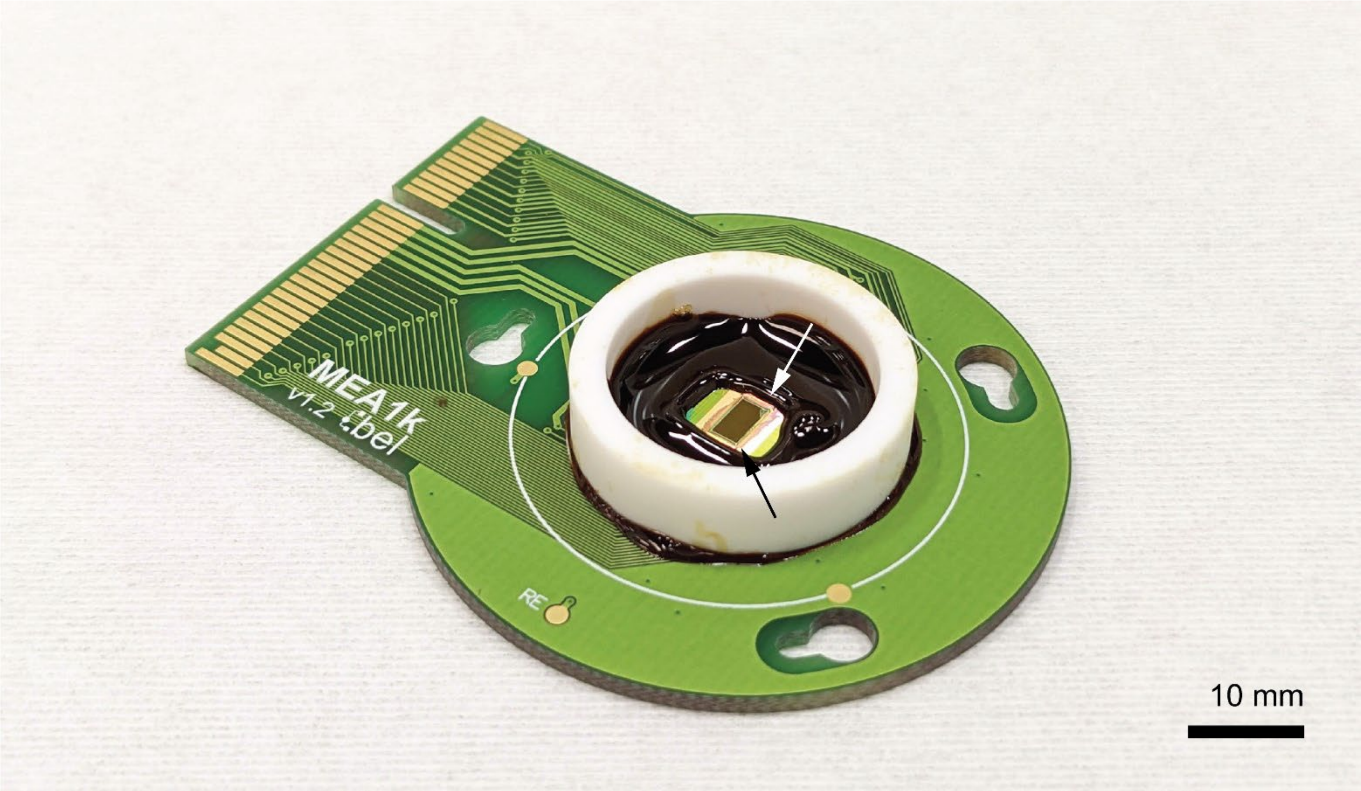

**Fig. S4 Packaged CMOS array**. The CMOS array printed with the microelectrodes were wire-bonded to a custom printed circuit board and the wire bonds were covered with epoxy. Arrows indicate locations where the microelectrodes were accidentally damaged during handling.

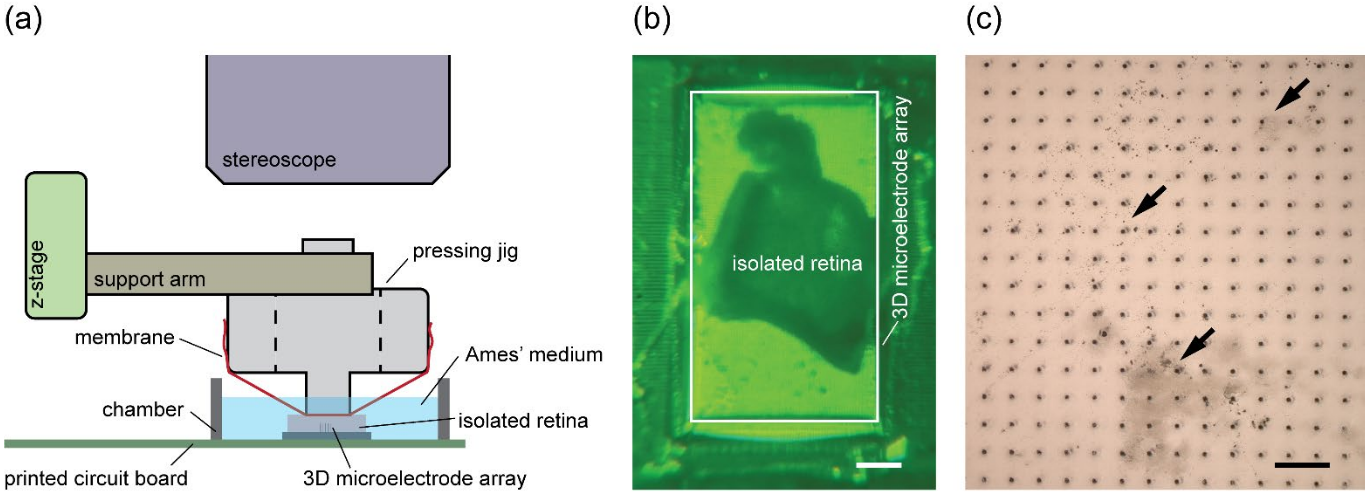

**Fig. S5 Retina pressing setup (a)** The pressing setup consists mainly of a *z*-stage, a support arm, and a custom-designed and 3D-printed pressing jig. The pressing jig has a through-hole in the middle (indicated by the dashed lines) that allows monitoring the tissue-pressing process when the *z*-stage is slowly lowered. A nylon mesh membrane is stretched taut across the tissue-facing end of the pressing jig to apply uniform pressure onto the tissue. **(b)** An image obtained with the stereoscope showing the isolated retina positioned on top of the 3D microelectrode array (outlined by the white rectangle) right before pressing. Scale bar: 500 μm. **(c)** An optical micrograph showing the microelectrode array remains largely intact after insertion into and removal from the retina. Residual retinal tissue (indicated by arrows) can be seen across the array. Scale bar: 50 μm.
