## Supplementary figures and images for "Direct-print three-dimensional electrodes for large-scale, high-density, and customizable neural interfaces"

### Supplementary Video

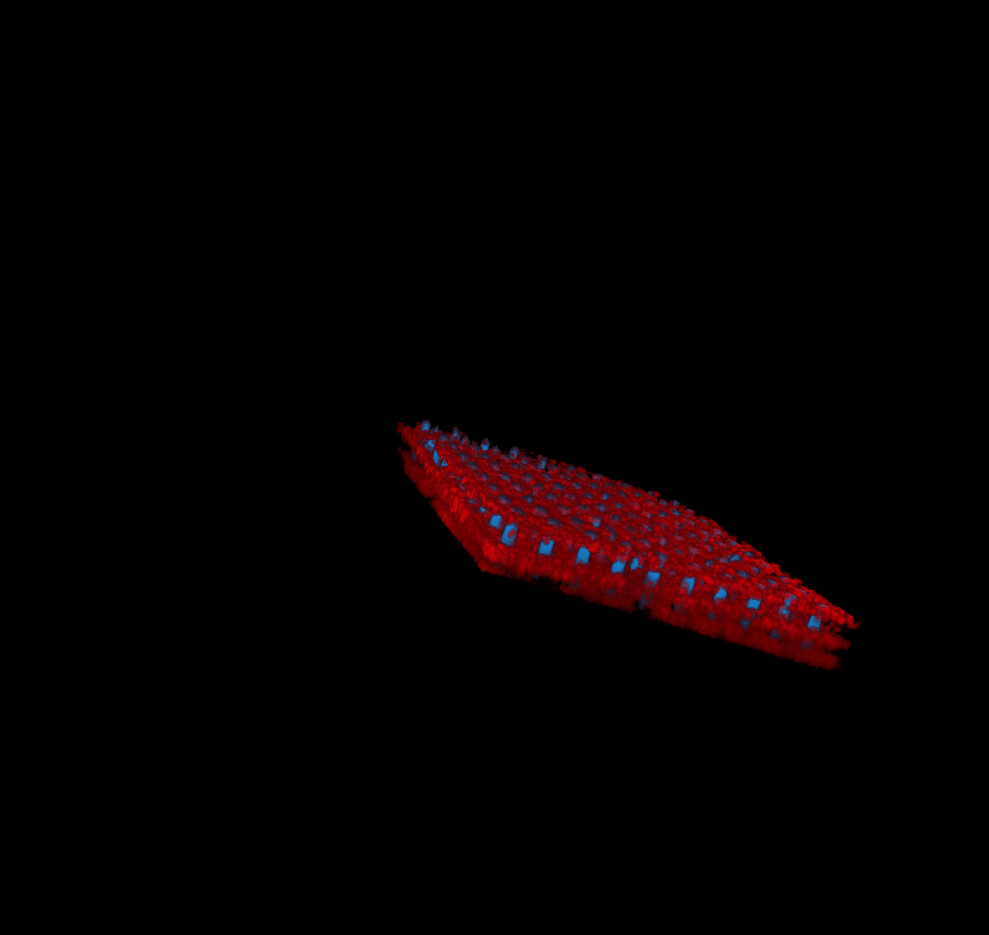
